# Revisiting Alpha Resting State Dynamics Underlying Hallucinatory Vulnerability: Insights from Hidden Semi-Markov Modeling

**DOI:** 10.1101/2023.12.11.571070

**Authors:** H. Honcamp, M. Schwartze, M. Amorim, D.E.J. Linden, A.P. Pinheiro, S.A. Kotz

## Abstract

Resting state (RS) brain activity is inherently non-stationary. Hidden Semi-Markov Models (HsMM) can characterize the continuous RS data as a sequence of recurring and distinct brain states along with their spatio-temporal dynamics. Recent explorations suggest that EEG brain state dynamics in the alpha frequency link to auditory hallucination proneness (HP) in non-clinical individuals. The present study aims to replicate these findings to elucidate robust neural correlates of hallucinatory vulnerability. Specifically, we aimed to investigate the reproducibility of HsMM states across different data sets and within-data set variants as well as the replicability of the association between alpha brain state dynamics and HP. We found that most brain states are reproducible in different data sets, confirming that the HsMM characterized robust and generalizable EEG RS dynamics. Brain state topographies and temporal dynamics of different within-data set variants showed substantial similarities and were robust against reduced data length and number of electrodes. However, the association with HP was not directly reproducible across data sets. These results indicate that the sensitivity of brain state dynamics to capture individual variability in HP may depend on the data recording characteristics and individual variability in RS cognition, such as mind wandering. We suggest that the order in which eyes-open and eyes-closed RS data are acquired directly influences an individual’s attentional state and generation of spontaneous thoughts, and thereby might mediate the link to hallucinatory vulnerability.

## 1. Introduction

### 1.1 Resting state dynamics

The resting brain transitions between episodes of engaged and disengaged functional networks. This is reflected in fluctuating neural oscillatory signatures on a sub-second timescale (Hutchison et al., 2013). Recent evidence suggests that continuous resting state (RS) activity can be systematically differentiated by a dynamic state allocation framework, in which the neural time series is segmented into a finite set of distinct but recurrent patterns of functional connectivity (FC), called brain states (Trujillo-Barreto, Araya, & El-Deredy, 2019; Vidaurre, Quinn, Baker, Dupret, Tejero-Cantero, & Woolrich, 2016; Woolrich et al., 2013). These brain states conform to underlying large-scale neural network activation patterns, and their switching dynamics relate to cognitive (dys-)function in healthy and patient populations (Kottaram et al., 2019; Nishida et al., 2013; Vidaurre et al., 2016). However, the states are “hidden”, as they cannot be directly observed.

### 1.2 HsMM for dynamic brain state allocation

Different methods have previously been used to identify brain states and to characterize their temporal dynamics, including the sliding window approach, microstate analysis, or several generative modeling approaches (Andreou et al., 2014; Geng, Xu, Sommer, Luo, Aleman, & Ćurčić-Blake, 2020; Kottaram et al., 2019; Trujillo-Barreto, Araya, & El-Deredy, 2019). As these methods differ considerably in their assumptions, advantages, and shortcomings, they are not equally suited to reveal the sub-second dynamics of the resting brain (H Honcamp, Schwartze, Linden, El-Deredy, & Kotz, 2022; Hutchison et al., 2013; Rukat, Baker, Quinn, & Woolrich, 2016).

Probabilistic generative methods, such as the Hidden Semi-Markov Model (HsMM), conceive the brain as a hybrid dynamic system, in which a fixed number of hidden states emit the continuous time series data that can be recorded with adequate neuroimaging methods such as electroencephalography (EEG) (Trujillo-Barreto, Araya, & El-Deredy, 2019). Unlike other approaches, the HsMM is not restricted by a pre-defined temporal window or a fixed number of states in its characterization of brain state dynamics (H Honcamp et al., 2022; Trujillo-Barreto, Araya, & El-Deredy, 2019). The HsMM also overcomes the main limitation of the classic Hidden Markov Model (HMM) by explicitly estimating state durations, which allows more accurate estimation of long-term dependencies of neural time series (Buzsáki & Mizuseki, 2014; Roberts, Boonstra, & Breakspear, 2015). Successful application of the HsMM to functional magnetic resonance imaging (fMRI) and magneto-/electroencephalogram (M/EEG) data showed that the temporal dynamics of brain states reflect between-subject variability in cognition and behavior (Honcamp et al., 2023; Kottaram et al., 2019; Vidaurre et al., 2016). The HsMM is thus a promising means for accurate and highly temporally resolved characterizations of early predictive neural markers of psychopathology (H Honcamp et al., 2022; Trujillo-Barreto, Araya, & El-Deredy, 2019).

### 1.3 Hallucinatory vulnerability

Following this rationale, recent work explored the applicability of a Bayesian formulation of the HsMM to assess whether temporal brain state dynamics in the alpha frequency range relate to inter-individual differences in hallucinatory predisposition in a non-clinical participant sample (Honcamp et al., 2023).

Hallucinatory experiences offer a particularly viable test case as they are essentially untriggered (i.e., spontaneous) sensory events in one or in different modalities that occur in the absence of a corresponding external source (Johns & Van Os, 2001). Evidence suggests that hallucinations result from dysfunctional source attribution, i.e., difficulties differentiating between internally and externally generated sensory events (Badcock & Hugdahl, 2012; Pinheiro, Farinha-Fernandes, Roberto, & Kotz, 2019). Although they are a cardinal symptom of psychotic disorders, most prominently schizophrenia, hallucinations are experienced by various other patient populations as well as by individuals from the general population (Bartels-Velthuis, Jenner, van de Willige, van Os, & Wiersma, 2010; Johns et al., 2014; Linszen et al., 2022). Accordingly, the psychosis continuum hypothesis suggests that the neural correlates of hallucinatory experiences in non-clinical individuals (i.e., without a formal diagnosis) are an attenuated version of those observed in clinically diagnosed individuals (Johns et al., 2014). The continuity of hallucinatory experiences is often referred to as hallucination proneness (HP), and is empirically supported by neuroimaging, electrophysiological, and behavioral studies (Allen, Freeman, Johns, & McGuire, 2006; Allen, Johns, Fu, Broome, Vythelingum, & McGuire, 2004; Badcock & Hugdahl, 2012).

Hallucinations and hallucinatory vulnerability are associated with several electrophysiological markers, including alterations in specific EEG frequency bands (Ford et al., 2012; Kindler, Hubl, Strik, Dierks, & Koenig, 2011). For example, alpha (8-12 Hz) power changes are linked to the perception threshold for sensory information, thus reflect individual variability in sensory sensitivity (Craddock, Poliakoff, El-Deredy, Klepousniotou, & Lloyd, 2017; Ecsy, Jones, & Brown, 2017; Shen, Han, Chen, & Chen, 2019). Hence, alpha band activity might indicate whether sensory events are (accurately) detected and therefore might also play a role in auditory phantom perceptions. RS alpha power fluctuations have been linked to changes in cognitive control (Clements, Bowie, Gyurkovics, Low, Fabiani, & Gratton, 2021). Lastly, alpha power also correlates with changes in attentional focus, e.g., during mind wandering and attentional switching between the external world and the internal physiological state (da Silva, Gonçalves, Branco, & Postma, 2022; Webster & Ro, 2020). Alpha band activity thus seems to relate to the brain’s dynamic adaptation to the variable demands of distinct attentional states. In turn, increased hallucinatory vulnerability is characterized by pronounced alterations in perceptual sensitivity, cognitive control, and attentional focus (Paulik, Badcock, & Maybery, 2008; Perona-Garcelán et al., 2014; F. Waters et al., 2012).

RS dynamics in the alpha band reflect individual differences in the vulnerability to auditory hallucinatory experiences (Honcamp et al., 2023). Specifically, mean activation duration and relative time spent in a distinct brain state characterized by auditory, somatosensory, and posterior default-mode network activity positively correlated with auditory (verbal) hallucinatory vulnerability. These findings suggest that alpha state dynamics during rest may reflect individual variability in attentional bias towards internal sensory events and altered perceptual sensitivity in the auditory domain. However, this study was the first investigation into the predictive value of HsMM state dynamics for HP and it involved a relatively small group of individuals. This warrants further systematic assessment of the reliability of the approach and the sensitivity of dynamic features of continuous EEG RS data as clinically relevant risk markers.

Considering data-inherent uncertainty (i.e., different sources of noise) and the stochastic nature of computational modeling (Beam, Manrai, & Ghassemi, 2020; Miłkowski, Hensel, & Hohol, 2018), the current study investigated (i) whether HsMM brain states are reproducible across different data sets and (ii) whether temporal dynamics of a given state are reproducible and robust predictors of non-clinical HP. Verifying the robustness of analysis tools and the replicability of research findings is indispensable to ensure that the identified electrophysiological markers of cognition and behavior are accurate and relevant. As the HsMM is still a novel tool to characterize brain state dynamics, the replication of findings also contributes to the identification of brain states that are robust to differences in recording procedures, number of electrodes, and other external influences. Consequently, discovering replicable brain states in different data sets further aids the physiological interpretation of the states, which will ultimately facilitate a better understanding of the underlying mechanisms of the HP continuum. To this end, data recorded and analyzed in Honcamp et al. (2023) was used as a reference data set and compared to a newly acquired replication data set. As this replication experiment is exploratory in nature, no specific hypotheses were generated.

## 2. Methods

Two data sets were used for the current replication study. Data set 1 (DS1) was obtained in our previous study (Honcamp et al., 2023) and consisted of 33 individuals varying in HP. Details on the sample characteristics, data collection, and processing can be found in Honcamp et al. (2023). Data set 2 (DS2) was obtained at the University of Lisbon, Portugal. Relevant details regarding participant recruitment and data processing are reported below. To maximize comparability to the results obtained in DS1, the data (pre-)processing, analysis, and modeling pipeline was adapted from Honcamp et al. (2023).

### 2.1 Participants and procedure

Ethical approval for DS2 was granted by the Deontological Committee of Faculty of Psychology at the University of Lisbon. Inclusion criteria for participation were: 1) native European Portuguese speaker; 2) right-handedness; 3) no history of psychiatric, neurological, or major medical illness, or clinical diagnosis of drug or alcohol abuse; 4) no present medication for clinical disorders that would affect EEG morphology, and 5) normal or corrected-to-normal vision and hearing. The final dataset (DS2) consisted of 65 individuals from the general population (50 females, 15 males, mean age = 21.57, SD = 6.593, age range = 18-48). Participation was compensated by course credits or a 10 € voucher. Participants provided written informed consent prior to participation.

Data collection took place at the University of Lisbon. Participants were invited for a behavioral assessment and an EEG recording session. HP was assessed using a validated Portuguese version of the Launay-Slade Hallucination Scale (LSHS) (Castiajo & Pinheiro, 2017). The LSHS consists of 16 items and specifically probes hallucinatory predisposition in non-clinical individuals. The items are scored on a 5-point Likert scale ranging from 0 (“certainly does not apply to me”) to 4 (“certainly applies to me”). Higher scores indicate a higher vulnerability to hallucinatory experiences. For each participant, we obtained the total HP score, reflecting an individual’s general predisposition to hallucinatory experiences as well as scores of the 5-item auditory HP (A-HP) and the 3-item auditory verbal HP (AV-HP) subscales (Honcamp et al., 2023; Pinheiro, Schwartze, Amorim, Coentre, Levy, & Kotz, 2020).

### 2.2 EEG data recording and processing

The EEG recording took place on the same day after the behavioral assessment. RS EEG data were recorded for 6 minutes, split into 3 minutes eyes-closed (EC) and 3 minutes eyes-open (EO) in the same order for all participants using a 64-channel Active Two BioSemi system (https://www.biosemi.com/products.htm). EEG recordings took place in an acoustically and electrically shielded EEG booth. Prior to the start of the recording, participants were asked to minimize bodily movements (including facial movements and blinking) and to stay awake. Between the EC and EO conditions, the experimenter entered the booth and instructed the participants explicitly to open their eyes.

EEG data were preprocessed using EEGLAB v2021 (Delorme & Makeig, 2004). As the length of the raw EC and EO data segments varied between participants, all data segments were cut to the same length of 3 minutes. Like in Honcamp et al. (2023), only EC data were used in the current replication. Data preprocessing included downsampling to 512 Hz, band-pass filtering between 1 and 40 Hz, rejection and spline interpolation of noisy channels using the clean_rawdata algorithm of EEGLAB (flat-line criterion = 5; channel criterion = 0.8; line noise criterion = 4; mean number of rejected and interpolated channels per participant = 5.17; SD = 3.13), average re-referencing, Automatic Subspace Reconstruction (ASR) with a burst criterion of 20 as implemented in EEGLAB, and combined ICA/PCA for identification of noisy components and control for rank deficiency. Components corresponding to eyeblinks, horizontal and vertical eye movements, remaining noisy channel activity, muscle-related activity, and line noise were removed (mean = 5.92; SD = 2.00).

### 2.3 Generation of data set variants

To match the sample size of DS1, DS2 was randomly divided into two smaller data subsets with 33 and 32 participants, respectively, from now on referred to as DS2a and DS2b. This ensured that any state differences between DS1 and DS2 were not due to discrepant sample size and any observed differences between DS2a and DS2b could not be attributed to differences in recording environment, hardware, number of electrodes, or recording time. Additionally, to account for variability in LSHS scores between DS1 and DS2, we selected another data subset, referred to as DS2c, with HP, A-HP, and AV-HP scores roughly matching those of the participants in DS1. An overview of sample characteristics of all data sets and modifications is provided in Table 1.

**Table 1.**
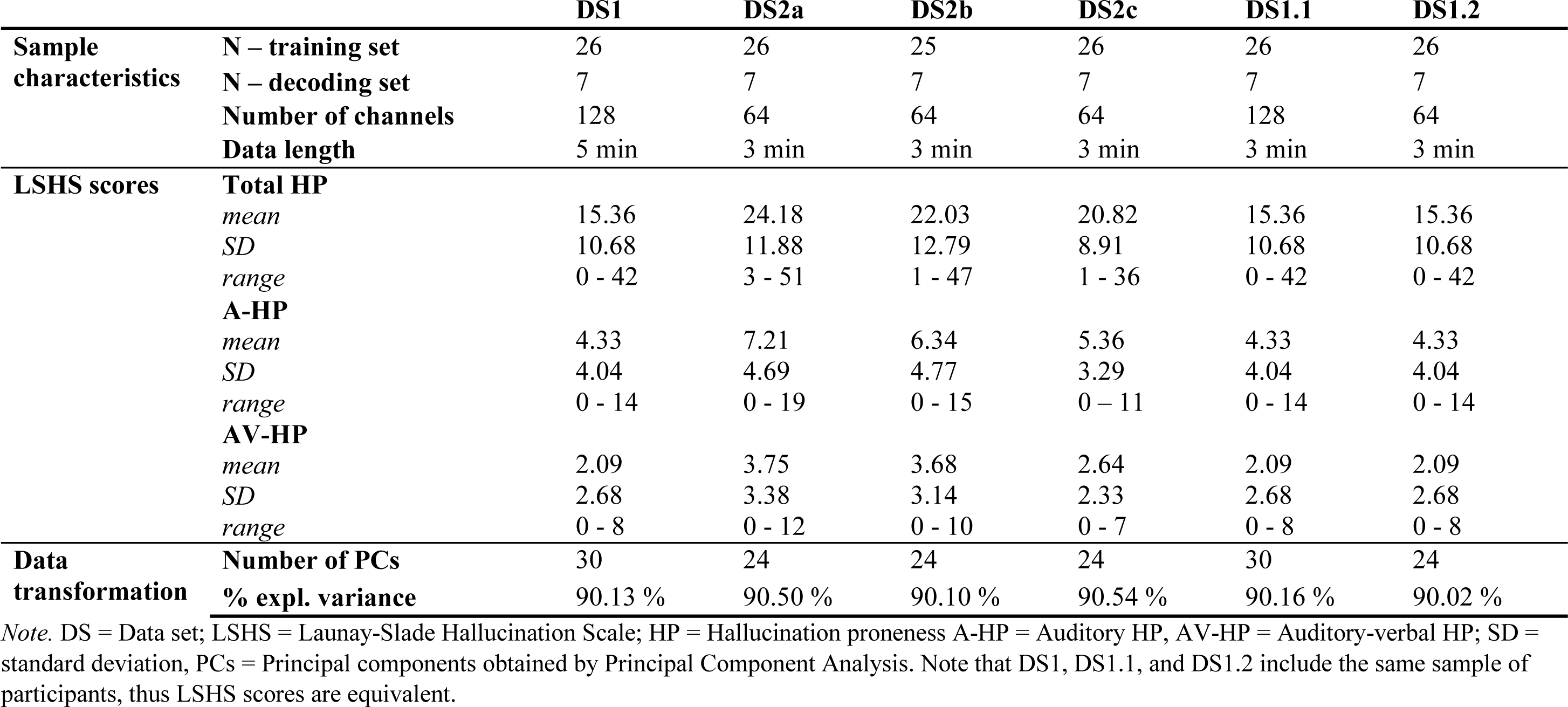
Descriptive statistics of included dataset variants.

### 2.4 Data transformation and feature extraction

Fig. 1 provides a schematic visualization of the data processing and analysis pipeline. After preprocessing, all DS2 variants were divided into two subsets: “normative” and “deviant”. The “normative” set comprised participants with the lower two-thirds of the HP scores, while participants with the upper third of HP scores served as the “deviant” set. This splitting prevented introducing bias in the estimation of brain state dynamics, i.e., conflating presumed normative and deviant dynamics of low and high hallucination-prone individuals, respectively.

**Fig. 1.**
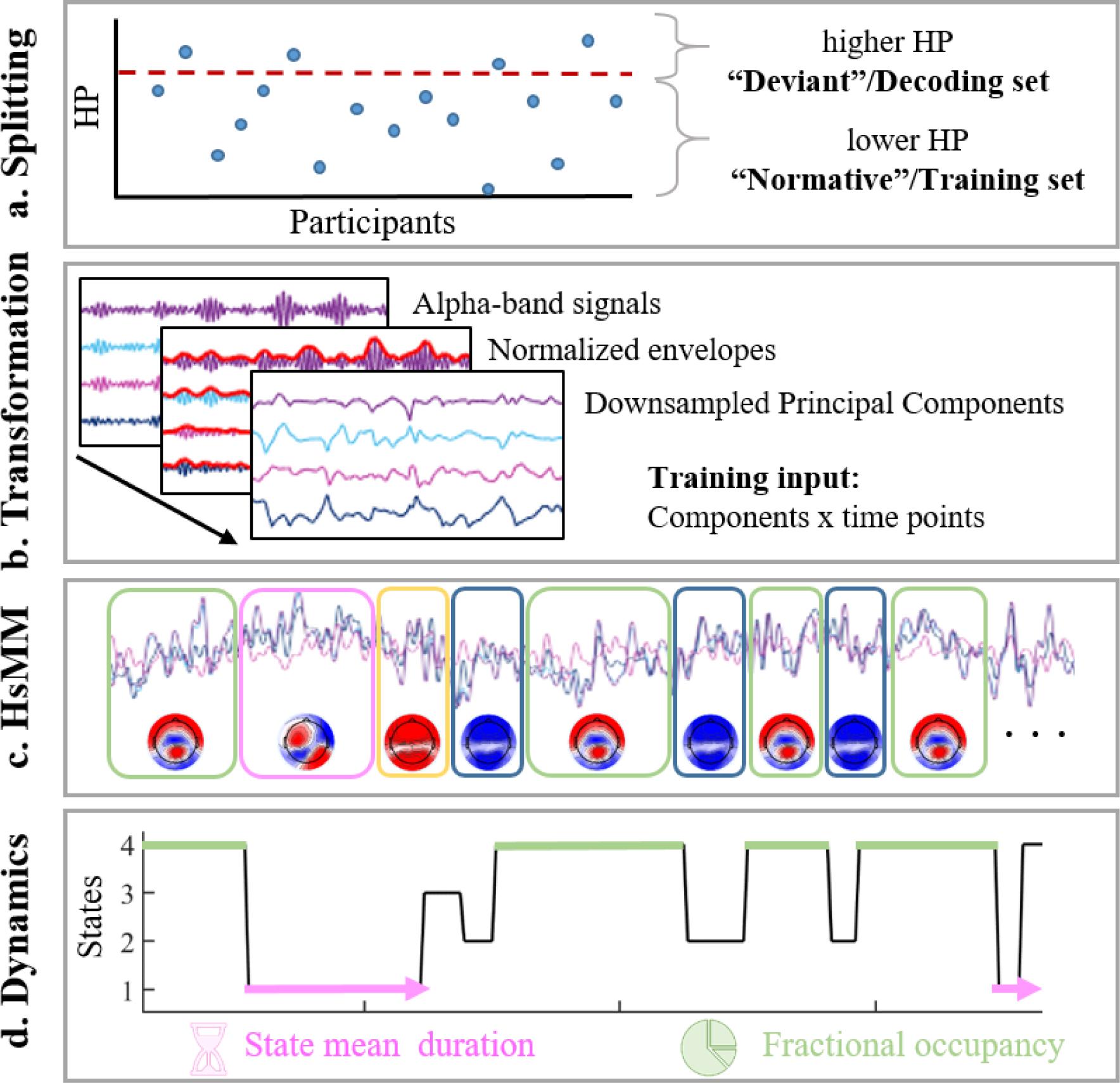
Schematic visualization of the data processing and analysis pipeline. **a. Splitting.** The schematic graph depicts the distribution of hallucination proneness (HP) scores of all participants within a data set (each point refers to one participant). Each individual data set was split into a training ("normative") and a decoding ("deviant") set, consisting of the lower two-thirds and the upper third of HP scores, respectively, illustrated by the dashed red line. **b. Transformation.** Data transformation included filtering to the alpha band (8-12 Hz), Hilbert transform, normalization, logarithmic transformation (not shown here), Principal Components Analysis (retaining >90% of explained variance of the data), and downsampling of the component data. The concatenated data (components x time points) of all participants were used as input to the model. **c. HsMM.** State allocation was performed by applying a Hidden semi-Markov Model (HsMM) to the concatenated data of all participants within a data set. **d. Dynamics.** The temporal brain state dynamics were calculated directly from the state sequence. The state’s mean duration is the average activation time of a state in seconds, while the fractional occupancy refers to the relative time occupied by a state in percent.

Data transformation and modeling were performed in MATLAB v2020b using the Brain Dynamics Toolbox and custom MATLAB scripts (Trujillo-Barreto, Araya, & El-Deredy, 2019). Each individual training set was then transformed separately according to the following steps. Data of the training sets were band-pass filtered between 8-12 Hz and detrended. A Hilbert transformation was performed to extract the signal envelope, which was then normalized for each participant by the global standard deviation across channels. Data were then logarithmically transformed. To increase computational efficiency, the data of all participants were temporally concatenated and subjected to a PCA. We reduced each data set to 24 principal components, retaining at least 90% of the explained variance (Table 1). Lastly, the PCA-reduced normalized envelope signal was downsampled to 64 Hz. The transformed data, concatenated for all participants (components x time points), then served as the input to the HsMM. The data of each respective “deviant” sub-set of all data sets were then transformed accordingly, using the PCA coefficients obtained from each respective training set. The latter ensured transforming the data into the same dimensional space.

### 2.5 Brain state allocation using HsMM and estimation of state dynamics

We applied a HsMM to each training data set separately to identify 5 transiently stable and recurrent brain states with distinct spatio-temporal characteristics. Preliminary data explorations showed that five states sufficiently characterized between-state variability (Honcamp et al., 2023). We used a Multivariate Normal (MVN) distribution to model the state emissions and a log-normal distribution to model the state durations. The variational inference algorithm was randomly initialized and repeated ten times to avoid dependence on initial conditions (Trujillo-Barreto, Araya, & El-Deredy, 2019).

The state dynamics of all individuals in each respective “normative”, i.e., training set, could be directly obtained from the resulting state sequence. State sequences of all individuals in the “deviant” sets were decoded by applying the already estimated model parameters from each corresponding training model output to the unseen data. Following this, two dynamic features were extracted for each state and each participant. Fractional Occupancy (FO), the total time occupied by a state compared to the total recording time, was calculated by summing up each individual activation duration divided by the total recording length of each respective data segment and is expressed as a percentage. The states’ mean duration (MD), the average time a state is active, was obtained by fitting a lognormal curve to the histogram of empirical state durations of each participant and state to extract the location (mu) and scale (sigma) parameters of the lognormal distribution. The lognormal mu values were then transformed to a normal scale to obtain interpretable duration values in milliseconds. Of note, duration values were only calculated if a participant visited a given state at least five times.

As the HsMM output corresponds to a PCA-reduced dimensional space, the state topographies cannot be directly interpreted. Therefore, state maps were obtained by projecting the mean of the estimated emission distributions from the HsMM back to the sensor space. To this end, the mean vector of the estimated MVN distribution for each state was multiplied by the PCA coefficients obtained during the data preparation.

### 2.6 Statistical analyses

Statistical analyses were performed in IBM Statistics SPSS v26 (IBM Corp., 2019) and MATLAB v2020b. To assess the reproducibility of the states’ characteristics (state maps and state dynamics) between data sets, we performed correlational analyses and Kolmogorov-Smirnov (KS) tests. The KS test quantifies the equality of continuous distributions of two samples according to the null hypothesis that the data from both samples were drawn from the same underlying distribution. To investigate the replicability of the relationship between the state dynamics (FO and MD) and HP within each sample, we performed non-parametric (Spearman’s Rho) correlations.

## 3. Results

### 3.1 Reproducibility of states between and within data sets

#### 3.1.1 State maps

The five HsMM state maps of DS1, DS2a, DS2b, and DS2c are depicted in Fig. 2, panel A. The order of the states as direct output of the HsMM is arbitrary. To aid visual comparison, we reordered the maps according to their closest match to the state maps obtained for DS1 as reported before (Honcamp et al., 2023). Overall, the state maps of all DS2 variants showed similarities with DS1 state maps. The most striking difference concerned the first state of DS1 (labeled state 1 in Honcamp et al., 2023), which was absent in DS2a, DS2b, and DS2c. However, the maps of DS2 variants were close to identical to each other, suggesting that the state maps can be robustly reproduced between different data set variants.

**Fig. 2.**
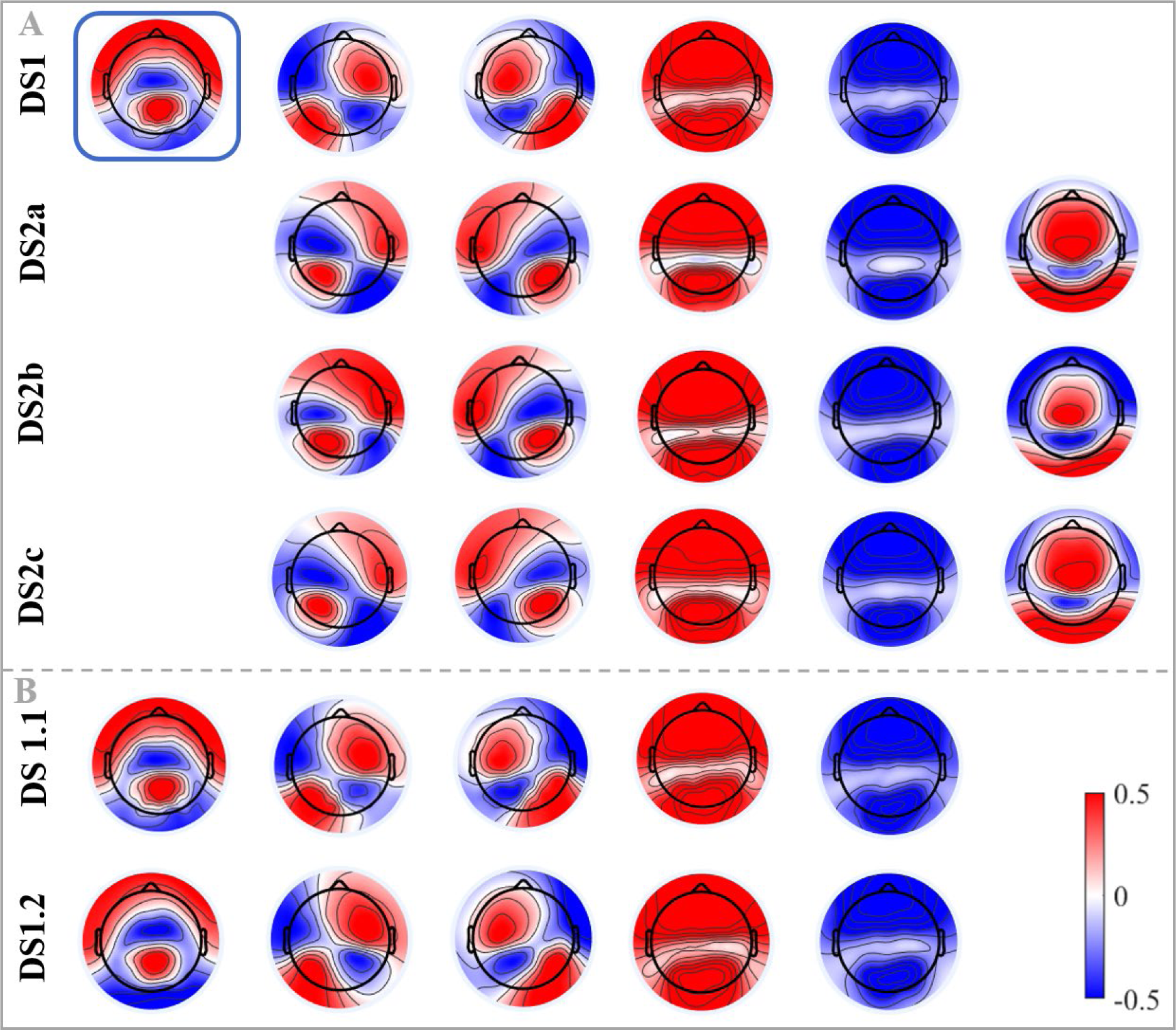
State maps of all data sub-sets. DS = Data set; **A. Between data set replication of state maps.** DS1 refers to the state maps obtained from 5 min EC RS data and 128 channels. The temporal dynamics of the highlighted state (blue rectangle) were found to be predictive of individual variability of auditory (verbal) hallucination proneness (Honcamp et a., 2023). DS2a refers to the state maps obtained from 3 min EC RS data and 64 channels corresponding to the first random sample of participants. Likewise, DS2b state maps correspond to the second random sample of participants. DS2c consists of a sample of participants with LSHS scores matched to those of DS1. Participants are different in DS1, DS2a, DS2b. Participants included in DS2c partly overlap with those of DS2a and DS2b. **B. Within data set replication of state maps.** DS1.1 refers to an adaptation of DS1.1 consisting of the first 3 min of the continuous EC data segment and all original 128 channels. Lastly, DS1.2 refers to the first 3 min of EC data from DS1 with 64 channels. The selected channels match those that are included in all DS2 variants. Participants of DS1, DS1.1, and DS1.2 are equal. See Table 1 for a comprehensive description of data sets.

As a follow-up analysis, we explored whether the difference in obtained state maps between DS1 and the DS2 variants resulted from i) the difference in data lengths (5 minutes DS1 versus 3 minutes DS2 variants) and ii) the additional difference in the number of channels (128 channels in DS1 versus 64 channels in DS2). To this end, we extracted the first 3 minutes from the already pre-processed data of DS1 and modeled the training data according to the same processing and training setting as described above. This adapted data set is referred to as DS1.1. Additionally, to explore the influence of the number of channels, we extracted 64 channels from DS1.1, corresponding to those used in all DS2 variants. The resulting data set, DS1.2, thus contained the first 3 minutes of EC RS data of 64 channels equivalent to DS2 variants. Data processing and transformation was conducted similarly to the steps reported above, except that we performed PCA with 24 (instead of 30) components to match the reduced data size and retain at least 90% of the explained variance. Moreover, data in DS1.2 was re-referenced to the common average again after extracting the 64-channel subset. The state maps obtained for DS1.1 and DS1.2 are shown in Fig. 2 panel B. Interestingly, state maps of all DS1 variants show substantial similarity, indicating the within-data set reproducibility of HsMM brain state maps. Consequently, the previously described differences between DS1 and DS2 are not attributable to the data recording length or the number of channels.

To quantify the similarity of state maps between DS1 and DS2 variants, we performed KS tests. Corresponding statistics can be found in Supplementary material A (Supplementary Tables 4 and 5). While the state maps of DS1 variants were not statistically different, the state maps of DS2 variants showed some differences, mainly concerning state 5, likely due to the extent of the fronto-central positivity (see Supplementary Fig. 1 and 2 for reference).

#### 3.1.2 State dynamics

We also investigated the consistency of empirical state duration and FO distributions across the different variants within data sets DS1 and DS2. Supplementary Fig. 3 and 4 (section A2) show the empirical state duration distributions for DS1 and DS2, respectively. The similarity between distributions was assessed by KS tests and Pearson’s R (Supplementary Table 6 and 7). For all comparisons of the same state across data set variants, the state’s duration distributions were not statistically different. Likewise, Supplementary Fig. 5 and 6 (section A3) show the states’ FO distributions for DS1 and DS2 variants, respectively. Supplementary Tables 8 and 9 depict the corresponding KS tests for equality of distributions. As for the state durations, the FO distributions per state were not statistically different across DS1 variants. Regarding the DS2 variants, only one state’s FO distribution was different between DS2a and DS2b.

### 3.2 Replicability of relationship between state dynamics and hallucination proneness

Honcamp et al. (2023) showed that FO and MD of one state (highlighted in blue in Fig. 2; referred to as state 1 in the original publication) were predictive of individual variability in auditory (and auditory verbal) hallucinatory vulnerability. However, the fact that this state was absent in all DS2 variants implies that this effect cannot be replicated. Exploratory non-parametric correlations between all states’ FO and MD values of DS2a, DS2b, and DS2c can be found in Supplementary Material B (Supplementary Tables 10 - 12).

To assess the replicability of the relationship between state dynamics and hallucinatory vulnerability in DS1 variants (DS1.1 and DS1.2), we performed non-parametric correlations between state 1 FO and MD values and general HP, A-HP, and AV-HP (Table 2). The correlational analysis showed a consistent pattern. The relationship between FO/MD and general HP was not significant in all DS1 variants, suggesting that none of the included metrics captured individual variability in general hallucinatory vulnerability as measured by the LSHS. Interestingly, in DS1.1, only FO was significantly associated with both A-HP and AV-HP, whereas in DS1.2, only MD showed a positive association. This discrepancy might reflect that different state dynamics are differently sensitive to data characteristics such as the data length or the number of electrodes.

**Table 2.** Non-parametric correlations (Spearman’s Rho) between state 1 fractional occupancy and mean duration values and hallucination proneness of DS1 variants.

|  |  | DS1 |  | DS1.1 |  | DS1.2 |  |
| --- | --- | --- | --- | --- | --- | --- | --- |
|  |  | FO | MD | FO | MD | FO | MD |
| **HP** | rho | .100 | .190 | .088 | .029 | .006 | .241 |
|  | p-value | .579 | .291 | .628 | .871 | .972 | .176 |
| **A-HP** | rho | .397 | .391 | .389 | .277 | .259 | .410 |
|  | p-value | .022* | .025* | .025* | .119 | .145 | .018* |
| **AV-HP** | rho | .408 | .459 | .390 | .333 | .140 | .408 |
|  | p-value | .019* | .007** | .025* | .058 | .437 | .018* |

*Note.* DS1 = Data set 1; 5 minutes, 128 channels, DS1.1 = Data set 1.1; 3 minutes, 128 channels; DS1.2 = Data set 1.2; 3 minutes, 64 channels; HP = Hallucination proneness A-HP = Auditory HP, AV-HP = Auditory-verbal HP, FO = fractional occupancy, MD = mean duration; Reported p-values are uncorrected; \*p<.05, \*\*p<.01; Reported p-values are uncorrected.

## 4. Discussion

The current study investigated (i) whether HsMM brain states in the RS alpha frequency band are reproducible across different data sets, and (ii) whether brain state temporal dynamics are robust predictors of non-clinical HP. To this end, we applied an HsMM to two independent data sets and within-data set variants to estimate five states per model along with their spatiotemporal characteristics. We further explored the reproducibility of state topographies, duration, and FO distributions between and within data sets.

We found that the five-states model solution was reproducible across different within data set variants and was robust against reduced data length and number of electrodes. Specifically, states of different variants of the same data set showed high degrees of spatial (i.e., state maps; Fig. 2) and dynamic (i.e., FO and duration distributions; Supplementary Fig. 3-6) similarity. The states further showed considerable overlap between data sets DS1 and DS2. Interestingly, the state (here, state 1 marked in blue in Fig. 2) that was associated with A-HP and AV-HP in DS1 (Honcamp et al., 2023) was not represented in either of the DS2 variants. This suggests that the two data sets were characterized by partially qualitatively different brain state topographies, despite controlling for the potential influence of data recording length (DS1.1), number of channels (DS1.2), and variability in LSHS scores (DS2c). Moreover, while the same state’s FO and MD values of DS1 variants showed a consistently positive association with A-HP and AH-HP (Table 2), the state dynamics across DS2 variants were rather randomly related to LSHS data (Supplementary Tables 10-12). In the following, we will discuss this finding, considering between and within-individual variability of RS FC and the influence of study design and data recording characteristics.

### 4.1 Temporal variability of the resting state

Time-averaged RS FC within large-scale neural networks (i.e., resting state networks, RSNs) shows high inter-individual consistency (Damoiseaux et al., 2006). However, there is considerable intra-individual variability in RSN connectivity and corresponding electrophysiological signatures across time (Baker et al., 2014; Hutchison et al., 2013; Woolrich et al., 2013). This neural variability reflects distinct cognitive modes, referred to as phenotypes of RS cognition, that characterize different stimulus-independent thought patterns and attentional states during unconstrained episodes of mind wandering (Diaz et al., 2014; Diaz et al., 2013; Gonzalez-Castillo, Kam, Hoy, & Bandettini, 2021). Using the Amsterdam Resting-State Questionnaire (ARSQ), Diaz et al. (2013) discovered seven dimensions of RS cognition, including Discontinuity of Mind, Self, and Somatic Awareness. The ARSQ-based dimensions not only revealed individual preferences for a given cognitive mode, but also linked to clinical measures and health status (e.g., depression, anxiety, and sleep quality). This is consistent with research on RS FC associated with cognitive-behavioral variability in healthy and patient populations (Vaidya & Gordon, 2013; Zhang et al., 2021). Therefore, a large proportion of variability in the estimated brain state dynamics might result from inter-individual differences in cognitive modes, or RS phenotypes. Moreover, RS dynamics in the Default-Model Network (DMN) are modulated by person-specific factors such as the circadian phenotype (Facer-Childs, Campos, Middleton, Skene, & Bagshaw, 2019), situation-specific factors, such as the amount (or lack) of external constraints, e.g., task instructions, (Christoff, Irving, Fox, Spreng, & Andrews-Hanna, 2016) as well as data recording characteristics (Van Diessen et al., 2015). The DMN is engaged in spontaneous thought, mind wandering, and self-referential processing (Raichle, 2015; Seeley et al., 2007). Mind wandering is defined as a mental state (or sequence of mental states) in which spontaneous, i.e., task- and stimulus-unrelated thoughts arise due to the absence of constraints on the content and the transition between different mental states, such as instructions of a highly demanding cognitive task that require externally focused attention (Christoff et al., 2016). Consequently, mind wandering naturally occurs when external events no longer capture an individual’s attention, i.e., during attentional decoupling and periods of low cognitive and perceptual load (Forster & Lavie, 2009; Schooler, Smallwood, Christoff, Handy, Reichle, & Sayette, 2011). Thus, inter-individual variability in RS cognition, attentional (de-)coupling, and DMN activation might partially explain the observed state differences between DS1 and DS2.

### 4.2 Resting state recording characteristics

FC and network-based metrics of RS M/EEG recordings are affected by several methodological choices in data recording and analyses (Gonzalez-Castillo et al., 2021; Van Diessen et al., 2015). For instance, the reliability of fMRI FC measures greatly improves with increased data length (Birn et al., 2013). Although we did not systematically investigate the effect of data length on the replicability of brain states here, we observed that the identified state maps remained highly stable when reducing the data set from 5 to 3 minutes (DS1 versus DS1.1). However, this does not exclude the possibility that a longer RS recording would yield a different set of brain states.

What seems critical in the current findings are the impact of condition, i.e., eye status, and condition order during the RS data recording. It has been consistently shown that brain state dynamics during rest qualitatively differ between open and closed conditions (Agcaoglu, Wilson, Wang, Stephen, & Calhoun, 2020; Agcaoglu, Wilson, Wang, Stephen, & Calhoun, 2019). Moreover, EO and EC conditions differently relate to vigilance and attentional state (Wong, DeYoung, & Liu, 2016) as well as to arousal, which strongly correlates with alpha power fluctuations (Barry, Clarke, Johnstone, Magee, & Rushby, 2007). Together, these findings suggest that network FC and corresponding electrophysiological signatures tend to be more reliable with longer recording duration and that they are modulated by the eye status during rest, which is directly related to an individual’s attentional status.

To overcome such condition-dependent differences, we opted to only include EC data in the current analyses. However, the RS recording of DS1 and DS2 differed in the order in which EO and EC data were obtained. Specifically, while the EC data in DS1 were recorded *after* 5 minutes of EO data had been collected, the EC data in DS2 were recorded for 3 minutes at the beginning of the recording session, i.e., *prior to* the recording of EO the data. This means that participants in DS1 were in the RS longer than participants in DS2, who did not do EO before, but were asked to close their eyes immediately at the beginning of the recording session. Given the time course of mind wandering and the generation of spontaneous thoughts, it is reasonable to assume that individuals in DS1 may have experienced attentional decoupling and started to mind wander earlier than the participants in DS2. This might be related to differences in external constraints (e.g., task instructions requiring externally focused attention), which are modulated by the time passing between receiving instructions and the actual start of the recording. This would suggest that underlying RSNs dynamics differ considerably not only depending on the RS eye status but also on the recording order and duration. In support of this hypothesis, a study comparing RS activity during different RS conditions showed that DMN connectivity significantly differed when EC data were recorded first from recordings where EC data was recorded after several other conditions, i.e., EO with and without a fixation cross (Yan et al., 2009). These findings also tentatively suggested that the amount of cognitive load, i.e., external constraints, during rest modulates mind wandering and daydreaming propensity.

In summary, large-scale network dynamics during rest link to attentional shifts from the external environment to internally oriented and spontaneous though processes, dependent on variability in external constraints (Christoff et al., 2016). Thus, during periods of low external constraints, externally generated sensory events might be more easily suppressed, which, in turn, facilitates internally focused cognition. This attentional decoupling might only manifest after an individual has been in a state of rest for some time. Therefore, it is plausible that the DS1 and DS2 data reflect fundamentally different episodes at rest, namely one in which individuals may still be attentive to their external environment (DS2), and one in which individuals have already decoupled from external constraints and shifted their attention to internal thought processes (DS1). The experience of (auditory verbal) hallucinations and heightened hallucinatory vulnerability have been related to deficient attribution of internally versus externally generated sensory events due to altered self- and source monitoring (Alderson-Day et al., 2014; Allen et al., 2006; Badcock & Hugdahl, 2012; Pinheiro et al., 2019). Further, non-clinical, high hallucination-prone individuals show higher self-focused attention as compared to individuals with lower HP (Perona-Garcelan et al., 2014). Therefore, distinct attentional states as described above may be differently sensitive to individual variability in hallucinatory vulnerability as measured at rest.

### 4.3 Limitations and future directions

#### 4.3.1 Number of states

The current study advances our understanding of HsMM-derived EEG RS brain dynamics across different data sets and their potential as a neural correlate of non-clinical hallucinatory vulnerability. However, it is limited by a number of methodological choices that can affect the interpretation of results and offer several possible venues for future research. In our original study (Honcamp et al., 2023), we opted for fixing the number of states to be identified by the HsMM to five, as this number sufficiently characterized between-subject differences in state dynamics. Following this decision, the replication experiments described here were also based on HsMMs with five states. Fixing the number of states is computationally efficient, however, it may not provide the best fitting model solution for the input data, which may also partly explain why one state (labeled state 1 in DS1 variants; see Fig. 2) was not represented in DS2. Previous research on the classic Hidden Markov Model applied to M/EEG data considered models with up to 12 states (Hunyadi, Woolrich, Quinn, Vidaurre, & De Vos, 2019; Kottaram et al., 2019; Vidaurre, Smith, & Woolrich, 2017). The choice for a given number of states can be based on the model’s Free Energy (FE), a model fit metric considering both the complexity and accuracy of the model (Trujillo-Barreto, Araya, & El-Deredy, 2019). However, exploring a different number of states would go beyond the scope of the current study. Future investigations should therefore consider a higher number of states to achieve a more accurate, and potentially more generalizable, representation of electrophysiological dynamics during RS.

#### 4.3.2 Validity of the LSHS composite scores

We measured HP using the 16-item version of the LSHS, a validated and widely used self-report measure of hallucinatory predisposition in non-clinical samples (Larøi & Van Der Linden, 2005). Next to a measure of general HP, we also considered the auditory and auditory-verbal composite scores consisting of 5 and 3 items, respectively, following (Pinheiro, Schwartze, & Kotz, 2018). Previous research exploring the underlying factor structure of the LSHS in non-clinical and clinical samples, however, did not consistently find the auditory, nor auditory-verbal items as a dissociable factor. Instead, many studies found a factor that was best described by multisensory HP, including visual, tactile, olfactory, and auditory psychotic-like experiences (Aleman, Nieuwenstein, Böcker, & De Haan, 2001; Castiajo & Pinheiro, 2017, 2021; Vellante, Larøi, Cella, Raballo, Petretto, & Preti, 2012). Moreover, a factor combining items from auditory and visual domains was most effective in differentiating between controls from psychotic individuals (Siddi et al., 2018). This may suggest that modality-specific, i.e., auditory subscales may not sufficiently capture individual variability in subclinical hallucinatory experiences and corresponding electrophysiological changes.

Interestingly, many studies found a distinct factor characterized by vivid daydreams and thoughts (sometimes referred to as vivid mental events) explaining a considerable amount of variance in the HP scores (Aleman et al., 2001; Castiajo & Pinheiro, 2017; Levitan, Ward, Catts, & Hemsley, 1996; F. A. Waters, Badcock, & Maybery, 2003). Other studies found a factor comprised by items tapping into daydreaming propensity and auditory phantom percepts, such as “The sounds I hear in my daydreams are usually clear and distinct” and “In the past, I have had the experience of hearing a person’s voice and then found no one was there” (Preti, Bolton, & Van De Ville, 2017; Siddi et al., 2018; Vellante et al., 2012). Although the link between vivid daydreaming, mind wandering, and hallucinatory vulnerability is not straightforward, there is some support to consider such an association. For instance, patients with schizophrenia report more frequent episodes of mind wandering, which is associated with the severity of positive symptoms (e.g., hallucinations, delusions; Shin, Lee, Jung, Kim, Jang, & Kwon, 2015). Moreover, the proneness to visual hallucinations has been linked to higher ratings of vivid imagery, which may contribute to confusing externally and internally generated images (Aynsworth, Nemat, Collerton, Smailes, & Dudley, 2017). Lasty, it has been suggested that the tendency to misattribute internally generated sensory events to an external source (and consequently perceived as a hallucination) is strongly modulated by the vividness of the sensory experience (Fazekas, 2021). However, this hypothesis still lacks sufficient empirical evidence and should be investigated further.

In summary, if we assume that variability in vivid daydreaming and mind wandering during EEG data collection underlies the state difference in DS1 and DS2, scores on the A-HP and AH-HP subscales of the LSHS may not be differentiated enough to consistently link to HsMM brain state dynamics. This might also explain the inconsistent correlational patterns between the state dynamics of DS2 variants and HP presented in Supplementary Tables 10 – 12. Thus, future studies should systematically investigate this and account for individual variability in vivid thoughts and imagery as well as mind wandering and daydreaming propensity.

#### 4.3.3 Other electrophysiological correlates

Related to the previous point, the exploration of other EEG frequency bands might offer a complementary perspective on the neurocognitive mechanisms underlying HP. Although alpha fluctuations have been extensively discussed as a potential electrophysiological correlate of mind wandering (e.g., Compton, Gearinger, & Wild, 2019; Dhindsa et al., 2019; Webster & Ro, 2020), changes in beta and theta power, as well as in beta-theta ratio, have also been associated with mind wandering and task-related attention reduction (da Silva et al., 2022). Therefore, considering brain state dynamics across multiple frequency bands as a function of both mind wandering and hallucinatory vulnerability could be a target for future research. Lastly, given the above-summarized evidence, it is plausible that the order in which the EO and EC conditions are recorded has a substantial impact on the brain’s electrophysiology and, therefore, accounts for the observed differences in brain states between the two data sets. However, this should be systematically explored in future research to elucidate the effect of condition order on HsMM brain states and their link to attentional states and spontaneous thoughts.

## 5. Conclusion

In this study, we aimed to reproduce HsMM brain states of RS EEG data in the alpha-band across different data sets and to investigate the relationship between the states’ temporal dynamics and hallucinatory vulnerability. Brain states of different within-data set variants showed substantial similarities and were robust against reduced data length and number of electrodes. Moreover, most brain states were reproducible in different data sets. However, we could not reproduce a previously identified link between state dynamics and HP. We suggest that the sensitivity of brain state dynamic features (such as fractional occupancy and mean duration) depends partly on the data recording characteristics and external constraints, which, in turn, can influence an individual’s attentional state. Thus, the relationship between HsMM brain state temporal dynamics and HP may be mediated by individual variability in attention focus (externally versus internally oriented) and the generation of spontaneous thoughts. Future research is required to systematically investigate this hypothesis.

## Conflict of interest statement

The authors declare no conflict of interest.

## Funding information

HH is funded by the Studienstiftung des Deutschen Volkes; SK is funded by the BIAL foundation [BIAL 102/2022]. AP is funded by the BIAL foundation [BIAL 146/2020]; MA is funded by Fundação para a Ciência e a Tecnologia [SFRH/BD/132170/2017].

## Ethics approval statement

Ethical approval was granted by the dutch Medisch-ethische toestingscommissie (METC 20-035; Dutch trial register no. 23432) (reference data set, DS1) and by the Deontological Committee of Faculty of Psychology at the University of Lisbon (DS2).

## CRediT author statement

HH: Conceptualization, Methodology, Software, Formal analysis, Data Curation, Writing - Original Draft, Visualization; MS: Conceptualization, Supervision, Writing – Review & Editing; MA: Resources, Writing – Review & Editing; AP: Resources, Writing – Review & Editing; AP: Resources, Writing – Review & Editing; DL: Writing – Review & Editing; SK: Conceptualization, Supervision, Resources, Writing – Review & Editing

## Acknowledgements

We would like to acknowledge Survarnalata Xanthate Duggirala for her support during data acquisition.

## Data and code availability

The software used for HsMM implementation in this manuscript is available at https://github.com/daraya78/BSD. Anonymized data can be shared with other researchers upon reasonable request to the corresponding author.

## Notes

### Competing Interest Statement

The authors have declared no competing interest.

