## Supplementary Material for "Revisiting Alpha Resting State Dynamics Underlying Hallucinatory Vulnerability: Insights from Hidden Semi-Markov Modeling"

### A. Comparison of brain state spatial and dynamic features of all data sets

This section contains information on the statistical comparison between data sets.

#### A1. Statemaps

This section contains the comparison of the state maps among data sets variants. The state maps were obtained by projecting the mean of the estimated HsMM emission distributions back to the sensor space, i.e., multiplying the PCA coefficients obtained during data transformation with the mean vector of the estimated MVN distribution for each state.

**Supplementary Figure 1. DS1 state maps for reference.**

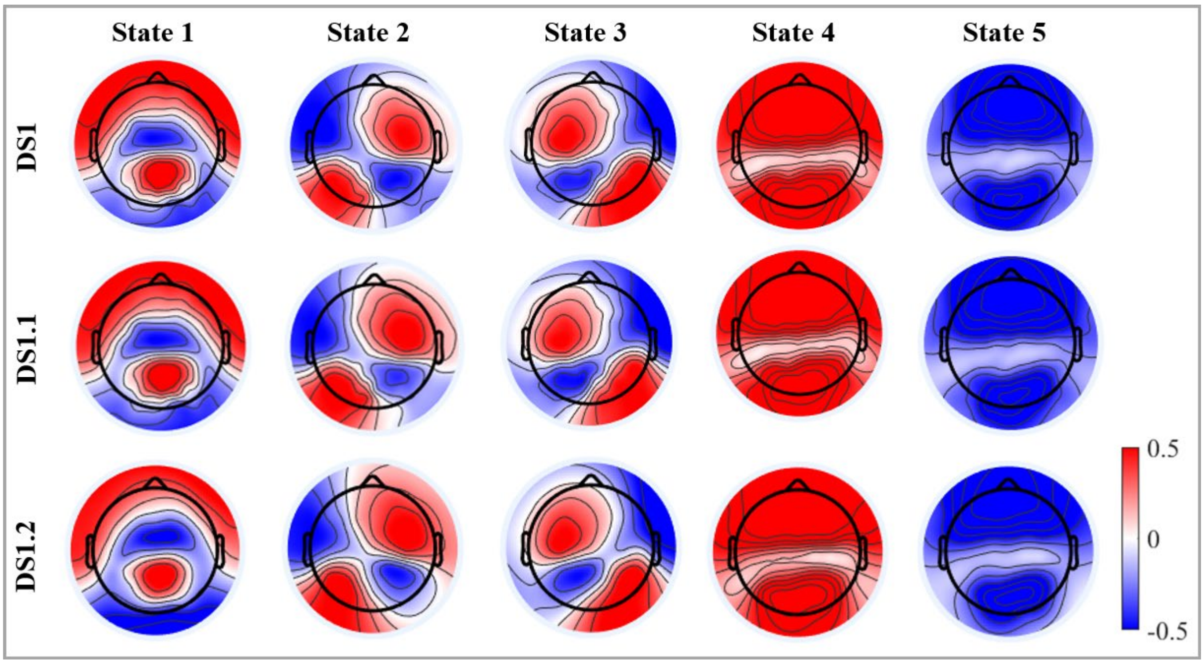


*Note.* States of DS1.1 and DS1.2 were ordered according to DS1 for statistical comparison.

**Supplementary Table 4. Comparison of statemaps between data set 1 variants**

| | | State 1 | State 2 | State 3 | State 4 | State 5 |
|---|---|---|---|---|---|---|
| **DS1 vs. DS1.1** | *KS statistic* | .055 | .102 | .063 | .102 | .047 |
| | *KS p-value* | .989 | .502 | .958 | .502 | .999 |
| **DS1 vs. DS1.2** | *KS statistic* | .125 | .156 | .063 | .125 | .133 |
| | *KS p-value* | .491 | .228 | .995 | .491 | .413 |
| **DS1.1 vs. DS1.2** | *KS statistic* | .109 | .086 | .063 | .141 | .094 |
| | *KS p-value* | .662 | .898 | .995 | .343 | .829 |

*Note.* KS = Kolmogorov Smirnov test; DS1 = Data set 1; 5 minutes, 128 channels, DS1.1 = Data set 1.1; 3 minutes, 128 channels; DS1.2 = Data set 1.2; 3 minutes, 64 channels; Pearson's R was omitted, as data sets consist of unequal number of entries (128 vs. 64). States were ordered according to DS1; see Supplementary Figure 1 for reference. Numbers are rounded to 3 decimal places.

**Supplementary Figure 2. DS2 state maps for reference.**

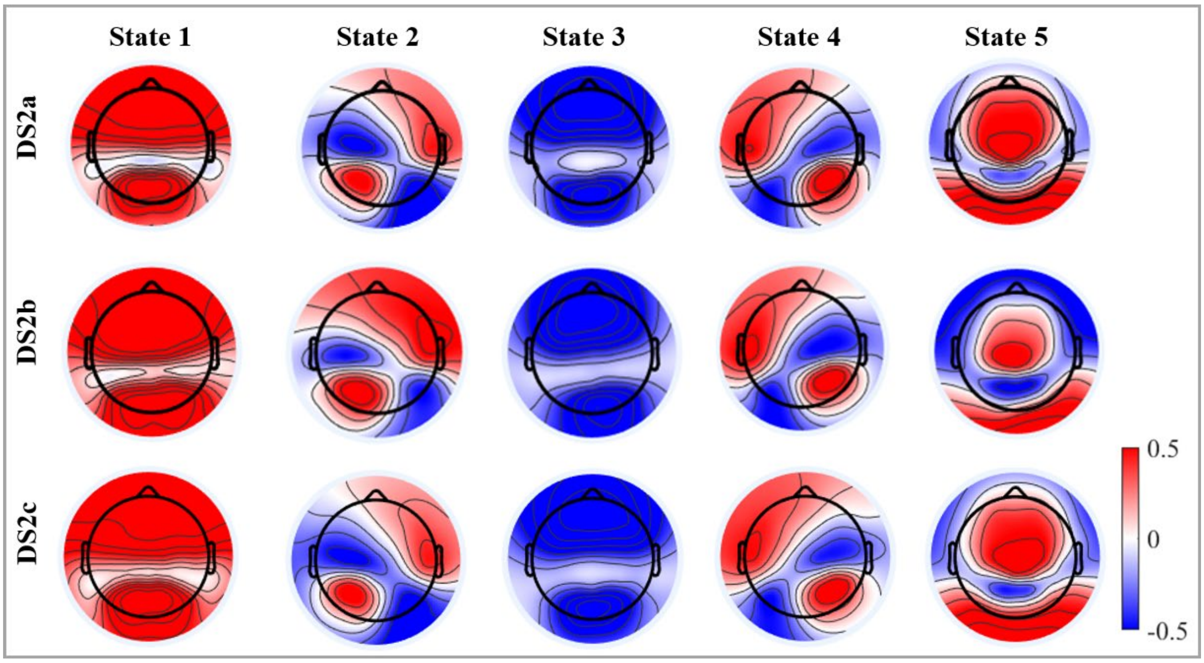


*Note.* States of DS2b and DS2c were ordered according to DS2a for statistical comparison.

**Supplementary Table 5. Comparison of statemaps between data set 2 variants**

| | | State 1 | State 2 | State 3 | State 4 | State 5 |
|---|---|---|---|---|---|---|
| **DS2a vs. DS2b** | *KS statistic* | .167 | .227 | .091 | .061 | .318* |
| | *KS p-value* | .291 | .056 | .937 | .999 | .0012 |
| **DS2a vs. DS2c** | *KS statistic* | .121 | .091 | .045 | .076 | .121 |
| | *KS p-value* | .689 | .937 | .999 | .989 | .689 |
| **DS2b vs. DS2c** | *KS statistic* | .152 | .258* | .091 | .106 | .288* |
| | *KS p-value* | .405 | .020* | .937 | .831 | .006* |

*Note.* KS = Kolmogorov Smirnov test; DS = Data set; DS2a = first random sample of DS2; DS2b = second random sample of DS2; DS2c = sample of DS2 with LSHS scores to DS1; *p < .05; States were ordered according to DS2a. See Table 1 for a comprehensive description of data sets; see Supplementary Figure 2 for reference. Numbers are rounded to 3 decimal places.

### A2. Empirical duration distribution

This section contains the comparison of the states' empirical duration distribution among data sets variants. The duration distributions depicted in Supplementary Figure 3 and 4 were generated by plotting a lognormal probability density function with the state's mu and sigma parameters for each data set variant.

**Supplementary Figure 3. Empirical duration distribution of DS1 variants**

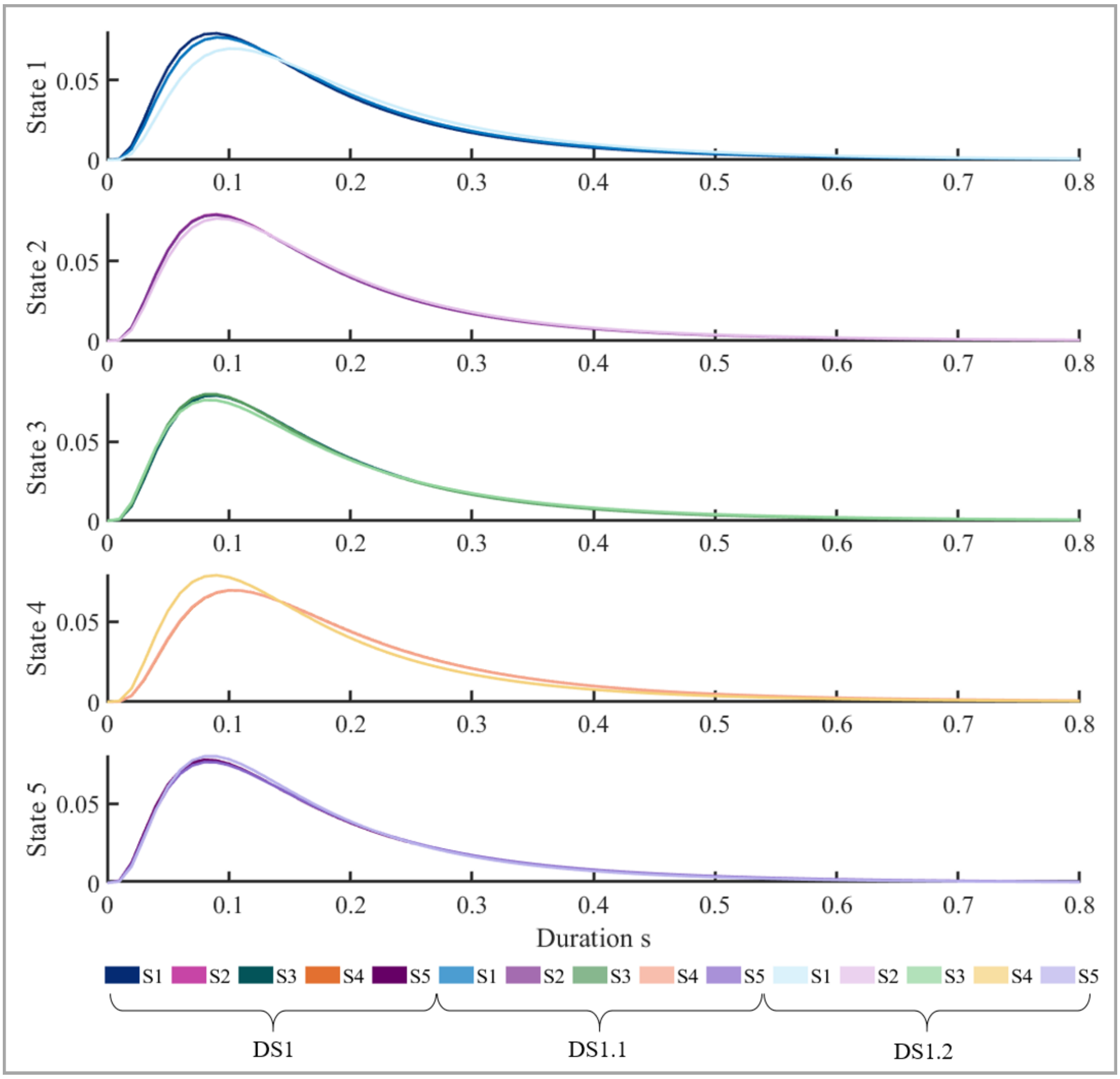


*Note.* DS = Data set; DS1 = 5 minutes, 128 channels, DS1.1 = 3 minutes, 128 channels; DS1.2 = 3 minutes, 64 channels. States were ordered according to DS1 state maps. See Table 1 for a comprehensive description of data sets.

**Supplementary Table 6. Comparison of empirical duration distributions between data set 1 variants**

| | | State 1 | State 2 | State 3 | State 4 | State 5 |
|---|---|---|---|---|---|---|
| **DS1 vs. DS1.1** | *KS statistic* | .029 | .013 | .023 | .016 | .010 |
| | *KS p-value* | .999 | 1 | 1 | 1 | 1 |
| | *Pearson's R* | 1 | 1 | 1 | 1 | 1 |
| **DS1 vs. DS1.2** | *KS statistic* | .042 | .016 | .058 | .039 | .055 |
| | *KS p-value* | .944 | 1 | .662 | .972 | .730 |
| | *Pearson's R* | .999 | 1 | .998 | .999 | .999 |
| **DS1.1 vs. DS1.2** | *KS statistic* | .029 | .023 | .051 | .039 | .051 |
| | *KS p-value* | .999 | 1 | .795 | .972 | .795 |
| | *Pearson's R* | 1 | 1 | .999 | .999 | .998 |

*Note.* KS = Kolmogorov Smirnov test; DS1 = Data set 1; 5 minutes, 128 channels, DS1.1 = Data set 1.1; 3 minutes, 128 channels; DS1.2 = Data set 1.2; 3 minutes, 64 channels. States were ordered according to DS1; see Supplementary Figure 1 for reference. Numbers are rounded to 3 decimal places.

**Supplementary Figure 4. Empirical duration distribution of DS2 variants**

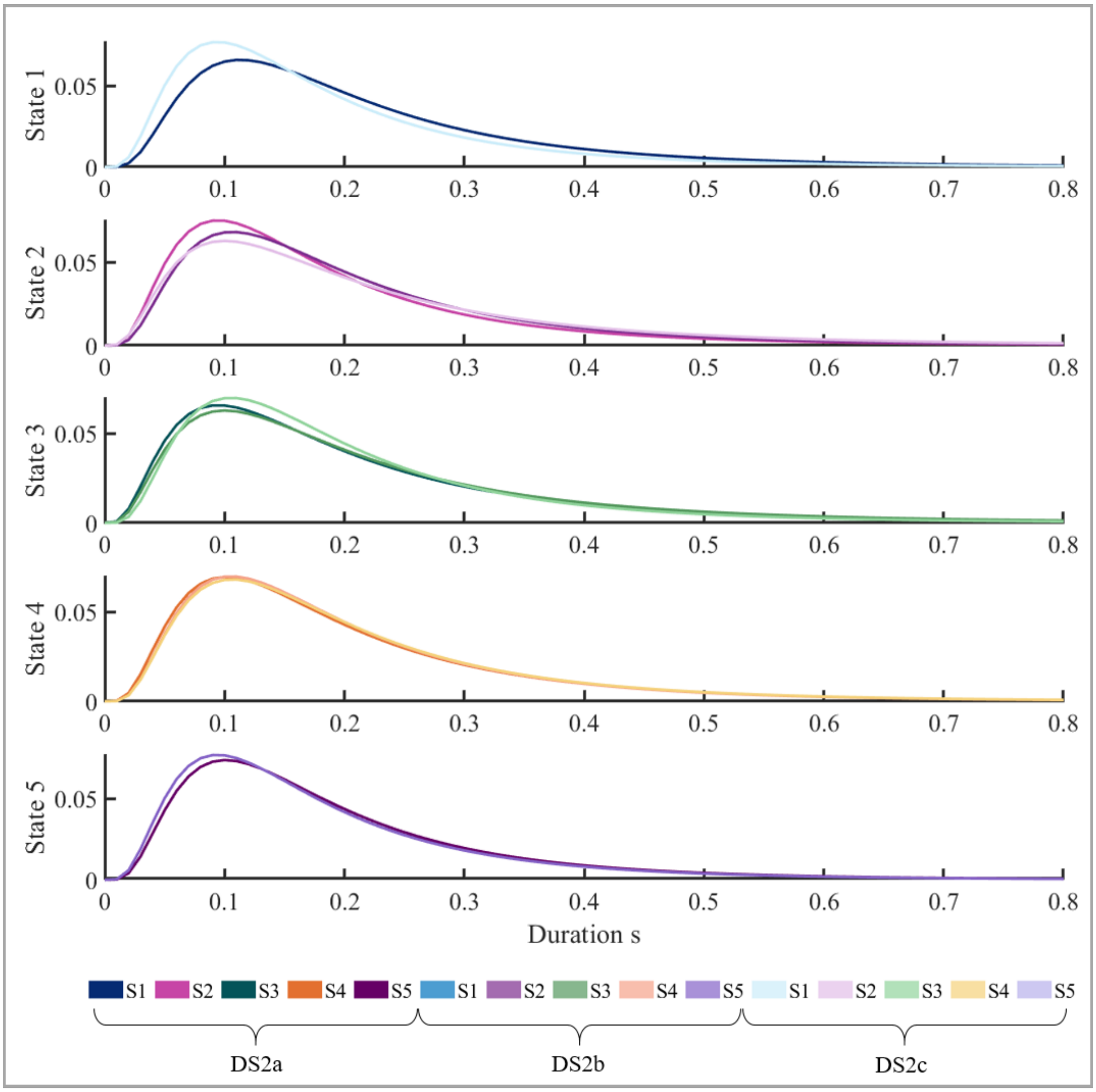


*Note.* DS = Data set; DS2a = first random sample of DS2; DS2b = second random sample of DS2; DS2c = sample of DS2 with LSHS scores to DS1. States were ordered according to DS2a state maps. See Table 1 for a comprehensive description of data sets.

**Supplementary Table 7. Comparison of empirical duration distributions between data set 2 variants**

| | | State 1 | State 2 | State 3 | State 4 | State 5 |
|---|---|---|---|---|---|---|
| **DS2a vs. DS2b** | *KS statistic* | .032 | .029 | .016 | .048 | .035 |
| | *KS p-value* | .997 | .999 | 1 | .854 | .989 |
| | *Pearson's R* | 1 | 1 | 1 | .999 | 1 |
| **DS2a vs. DS2c** | *KS statistic* | .032 | .077 | .090 | .039 | .032 |
| | *KS p-value* | .997 | .301 | .153 | .972 | .997 |
| | *Pearson's R* | 1 | .994 | .989 | 1 | 1 |
| **DS2b vs. DS2c** | *KS statistic* | .019 | .084 | .087 | .029 | .019 |
| | *KS p-value* | 1 | .217 | .183 | .999 | 1 |
| | *Pearson's R* | 1 | .992 | .990 | 1 | 1 |

*Note.* KS = Kolmogorov Smirnov test; DS = Data set; DS2a = first random sample of DS2; DS2b = second random sample of DS2; DS2c = sample of DS2 with LSHS scores to DS1. See Table 1 for a comprehensive description of data sets. States were ordered according to DS2a; see Supplementary Figure 2 for reference. Numbers are rounded to 3 decimal places.

### A3. Fractional Occupancy

This section contains the comparison of the states' fractional occupancy (FO) distribution among data sets variants. Regarding DS2 variants, note that DS2a and DS2c consisted of 33 participants, whereas DS2b consisted of 32 participants. For better visualization, the 33rd entry in DS2b was imputed by the mean FO value per state. However, for the statistical comparisons between DS2 variants (Supplementary Table 9), no imputed values were used.

**Supplementary Figure 5. Fractional occupancy distribution of DS1 variants**

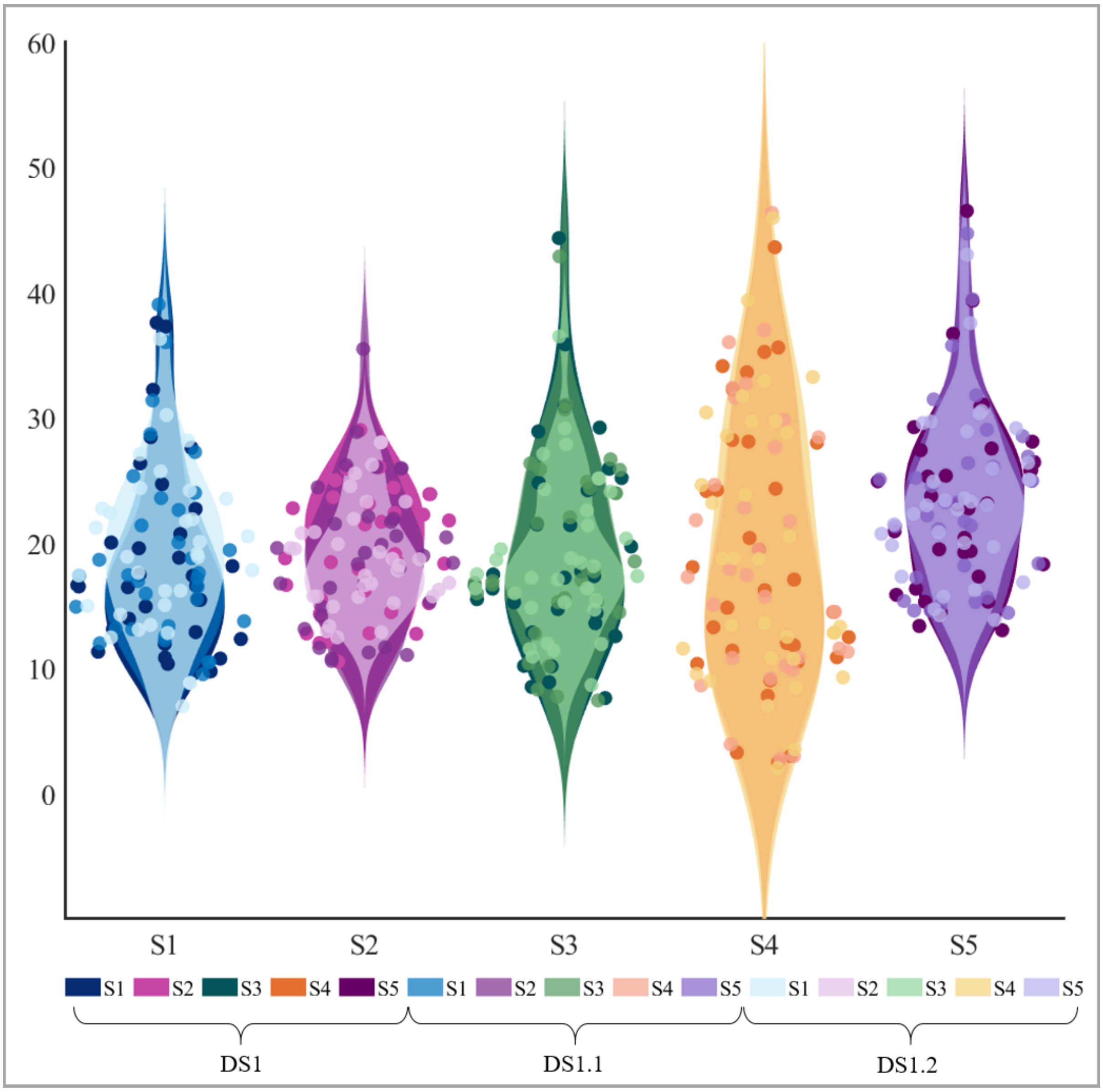


*Note.* DS = Data set; DS1 = 5 minutes, 128 channels, DS1.1 = 3 minutes, 128 channels; DS1.2 = 3 minutes, 64 channels. States were ordered according to DS1 state maps. Please refer to Table 1 for a comprehensive description of data sets.

**Supplementary Table 8. Comparison of fractional occupancy distributions between data set 1 variants**

| | | State 1 | State 2 | State 3 | State 4 | State 5 |
|---|---|---|---|---|---|---|
| **DS1 vs. DS1.1** | *KS statistic* | .121 | .152 | .121 | .091 | .121 |
| | *KS p-value* | .957 | .811 | .957 | .999 | .957 |
| **DS1 vs. DS1.2** | *KS statistic* | .212 | .212 | .121 | .121 | .152 |
| | *KS p-value* | .403 | .403 | .957 | .957 | .811 |
| **DS1.1 vs. DS1.2** | *KS statistic* | .182 | .182 | .121 | .121 | .121 |
| | *KS p-value* | .601 | .601 | .957 | .957 | .957 |

*Note.* KS = Kolmogorov Smirnov test; DS1 = Data set 1; 5 minutes, 128 channels, DS1.1 = Data set 1.1; 3 minutes, 128 channels; DS1.2 = Data set 1.2; 3 minutes, 64 channels; $^*p < .05$; see Table 1 for a comprehensive description of data sets. States were ordered according to DS1; see Supplementary Figure 1 for reference. Numbers are rounded to 3 decimal places.

**Supplementary Figure 6. Fractional Occupancy distribution of DS2 variants**

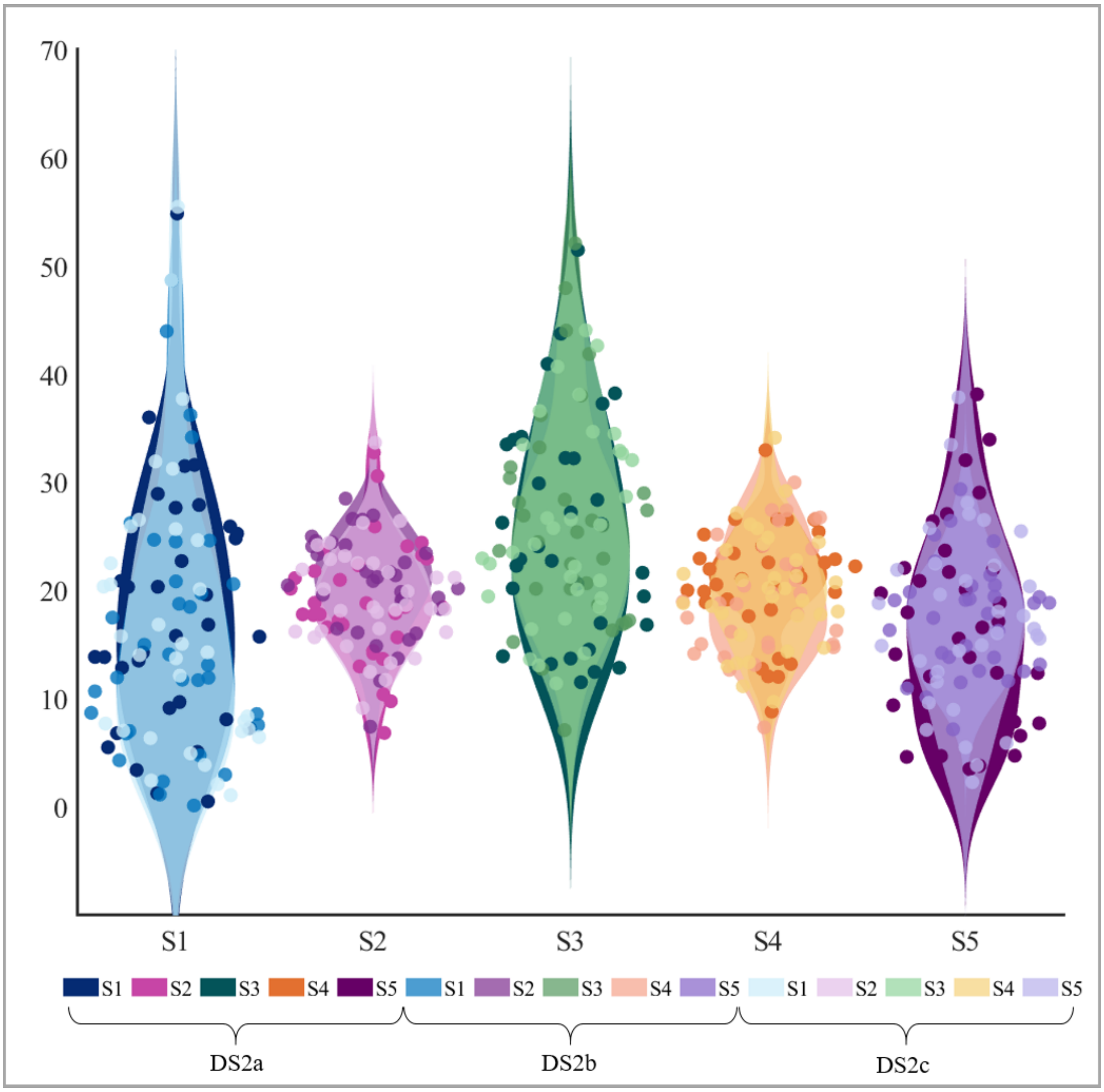


*Note.* DS = Data set; DS2a = first random sample of DS2; DS2b = second random sample of DS2; DS2c = sample of DS2 with LSHS scores to DS1. Please refer to Table 1 for a comprehensive description of data sets. Please note that DS2a and DS2c consisted of 33 participants, whereas DS2b consisted of 32 participants. For better visualization of the fractional occupancy (FO) distribution, the 33rd entry in DS2b was imputed by the mean FO value per state.

**Supplementary Table 9. Comparison of fractional occupancy distributions between data set 2 variants**

| | | State 1 | State 2 | State 3 | State 4 | State 5 |
|---|---|---|---|---|---|---|
| **DS2a vs. DS2b** | *KS statistic* | .198 | .356* | .150 | .291 | .241 |
| | *KS p-value* | .501 | .024* | .829 | .105 | .261 |
| **DS2a vs. DS2c** | *KS statistic* | .152 | .121 | .121 | .182 | .152 |
| | *KS p-value* | .811 | .957 | .957 | .601 | .811 |
| **DS2b vs. DS2c** | *KS statistic* | .113 | .295 | .115 | .164 | .181 |
| | *KS p-value* | .980 | .095 | .976 | .737 | .618 |

*Note.* KS = Kolmogorov Smirnov test; DS = Data set; DS2a = first random sample of DS2; DS2b = second random sample of DS2; DS2c = sample of DS2 with LSHS scores to DS1; *$p < .05$; see Table 1 for a comprehensive description of data sets. States were ordered according to DS2a; see Supplementary Figure 2 for reference. Numbers are rounded to 3 decimal places.

### B. Exploratory non-parametric correlations (Spearman's Rho) between state dynamics and hallucination proneness

This section contains exploratory non-parametric correlations between HsMM brain state dynamics and general hallucination proneness (HP), auditory HP (A-HP), and auditory-verbal HP (AV-HP), as measured by the Launay-Slade Hallucination Scale (LSHS), for all DS2 variants. The corresponding state maps can be found in Supplementary Figure 2.

**Supplementary Table 10.** Non-parametric correlations (Spearman's Rho) between state dynamics and hallucination proneness of DS2a

| | | **State 1** | | **State 2** | | **State 3** | | **State 4** | | **State 5** | |
|---|---|---|---|---|---|---|---|---|---|---|---|
| | | FO | MD | FO | MD | FO | MD | FO | MD | FO | MD |
| **HP** | *Rho* | .144 | .210 | .185 | .227 | -.057 | -.121 | -.202 | -.063 | -.002 | .019 |
| | *p* | .425 | .240 | .301 | .203 | .755 | .501 | .295 | .726 | .993 | .917 |
| **A-HP** | *Rho* | -.028 | .033 | .030 | .170 | .027 | -.067 | -.072 | .042 | .134 | .162 |
| | *p* | .879 | .856 | .869 | .344 | .884 | .711 | .698 | .817 | .456 | .367 |
| **AV-** | *Rho* | -.010 | .037 | -.037 | .217 | -.053 | -.081 | -.111 | .109 | .224 | .216 |
| **HP** | *p* | .955 | .339 | .838 | .225 | .769 | .652 | .537 | .547 | .210 | .228 |

*Note.* HP = Hallucination proneness A-HP = Auditory HP, AV-HP = Auditory-verbal HP, FO = fractional occupancy, MD = mean duration; Reported p-values are uncorrected; *p<.05, **p<.01; Reported p-values are uncorrected; Please refer to Table 1 for a comprehensive description of data sets. States were ordered according to DS2a; see Supplementary Figure 2 for reference.

**Supplementary Table 11.** Non-parametric correlations (Spearman's Rho) between state dynamics and hallucination proneness of DS2b.

| | | **State 1** | | **State 2** | | **State 3** | | **State 4** | | **State 5** | |
|---|---|---|---|---|---|---|---|---|---|---|---|
| | | FO | MD | FO | MD | FO | MD | FO | MD | FO | MD |
| **HP** | *Rho* | .319 | - | .038 | -.293 | .304 | -.096 | .270 | -.028 | .070 | -.149 |
| | *p* | .076 | .423*.<br>018 | .838 | .104 | .091 | .601 | .135 | .880 | .705 | .417 |
| **A-HP** | *Rho* | -.337 | - | -.059 | - | .355* | -.033 | .297 | -.021 | .042 | -.156 |
| | *p* | .059 | .421*.<br>013 | .747 | .405*.<br>022 | .046 | .859 | .098 | .909 | .819 | .395 |
| **AV-** | *Rho* | -.319 | - | -.200 | .459* | .298 | -.048 | .216 | -.013 | .329 | .086 |
| **HP** | *p* | .076 | .436*.<br>014 | .272 | .008 | .098 | .794 | .236 | .944 | .066 | .639 |

*Note.* HP = Hallucination proneness A-HP = Auditory HP, AV-HP = Auditory-verbal HP, FO = fractional occupancy, MD = mean duration; Reported p-values are uncorrected; *p<.05, **p<.01; Reported p-values are uncorrected; Please refer to Table 1 for a comprehensive description of data sets. States were ordered according to DS2a; see Supplementary Figure 2 for reference.

**Supplementary Table 12.** Non-parametric correlations (Spearman's Rho) between state dynamics and hallucination proneness of DS2c

| | | State 1 | | State 2 | | State 3 | | State 4 | | State 5 | |
|---|---|---|---|---|---|---|---|---|---|---|---|
| | | FO | MD | FO | MD | FO | MD | FO | MD | FO | MD |
| **HP** | *Rho* | -.109 | -.073 | .424* | .355* | .116 | .032 | -.065 | .038 | .119 | -.003 |
| | *p* | .546 | .687 | .014 | .043 | .521 | .861 | .719 | .853 | .510 | .986 |
| **A-HP** | *Rho* | -.306 | -.292 | .277 | .113 | .233 | -.014 | -.063 | -.009 | .319 | .163 |
| | *p* | .083 | .099 | .119 | .532 | .192 | .939 | .727 | .962 | .071 | .365 |
| **AV-HP** | *Rho* | - | - | .143 | .118 | .262 | .054 | -.023 | .074 | .441* | .328 |
| | *p* | .389*. 025 | .353*. 044 | .429 | .514 | .141 | .765 | .897 | .683 | .010 | .062 |

*Note.* HP = Hallucination proneness A-HP = Auditory HP, AV-HP = Auditory-verbal HP, FO = fractional occupancy, MD = mean duration; Reported p-values are uncorrected; *p<.05, **p<.01; Reported p-values are uncorrected; Please refer to Table 1 for a comprehensive description of data sets. States were ordered according to DS2a; see Supplementary Figure 2 for reference.
